# Meta-Analysis of EEG Findings on Pain Perception: Exploring Nociceptive and Neuropathic Pain Response Patterns

**DOI:** 10.1101/2023.10.31.564952

**Authors:** Lipnitskii Mikhail

## Abstract

This article presents a meta-analysis of research articles focusing on the use of electroencephalography (EEG) in the study of nociceptive and neuropathic pain perception. The objective of the study is to compare the findings of the reviewed articles with the three-route model of pain perception, which posits that specific brain regions are responsible for interpreting different aspects of pain. The articles included in the analysis were sourced from reputable databases such as Scopus, Google Scholar, and Pubmed. The selection criteria for these articles was based on the explicit demonstration of changes in EEG rhythms in response to pain sensations. This served as an important factor in determining their inclusion for further analysis. The results of the meta-analysis provide insights into the changes observed in EEG rhythms across different brain regions. By examining the location of these characteristic changes, the study makes assumptions about potential interrelationships between the observed EEG rhythms and the activity of specific brain regions discussed in the articles. Ultimately, this meta-analysis contributes to our understanding of the relationship between pain perception and EEG activity, shedding light on the potential role of distinct brain areas in processing different aspects of pain.

## 1 Introduction

Pain is a universal human experience and is one of the most common symptoms experienced by humans. It can be physical, emotional and psychological in nature and has a significant impact on people’s quality of life and well-being.

Understanding the mechanisms and processes involved in pain perception is essential for the development of effective treatments and management of pain. In recent years, research in the field of pain perception has gained new impetus for development. Thanks to advances in medical technology and neuroscience, scientists are able to study pain responses at the level of the brain and nervous system, allowing for a better understanding of how the brain perceives and processes pain signals and how these signals are transmitted throughout the body.

According to the nature of pain, a distinction is made between nociceptive and neuropathic pain[1].

Nociceptive pain refers to pain that arises from tissue damage and can manifest as either somatic or visceral pain. Somatic pain receptors are found in various structures such as the skin, subcutaneous tissue, fascia, connective tissues, periosteum, endosteum, and joint capsules. Stimulation of these receptors typically produces localized pain sensations, which can range from sharp to dull. However, burning pain is not commonly associated with involvement of the skin and subcutaneous tissue in the pathological process. On the other hand, visceral pain receptors are located in most internal organs and the surrounding connective tissue. When there is damage to the organ capsule or deep connective tissue elements, visceral pain may be more distinct and localized, often presenting as acute discomfort.

Neuropathic pain refers to a specific kind of pain that occurs due to abnormal activity of neurons in either the peripheral or central nervous system, which are responsible for responding to physical injury. This type of pain can be characterized by abnormal sensations, known as dysesthesia, or pain triggered by stimuli that would not typically cause pain, called allodynia. Neuropathic pain can manifest as a constant discomfort or occur in episodes, with the latter being described as sharp or resembling electric shocks.

Research on this process has been going on for some time, and there are currently several models that explain some aspects of the pain perception process:

- **Gating model of pain**. According to this model, pain signals are transmitted from damaged tissues through peripheral nerves to the spinal cord. In the spinal cord, pain signals pass through “gate cells” that act as a filter. Depending on various influences, the gate cells can either “open”, allowing pain signals to reach the brain and cause the sensation of pain, or “close”, blocking the transmission of signals and reducing the sensation of pain.
- **Model of the ascending pain system**. This model suggests that pain signals are transmitted from damaged tissues through nerve fibers to the brain, passing through various structures such as the spinal cord and thalamus. The brain further processes the signals, resulting in the sensation of pain.

Various electrophysiology methods are used to study brain processes. One such method is electroencephalography (EEG), which provides information about activity on the surface of the cerebral cortex. The process of pain perception has also been studied in detail using fMRI [2, 3, 4, 5, 9], CT[6, 7, 8], PET[10, 11, 12], but the most relevant method of research was the electroencephalography(EEG) method.

The purpose of this study is to compare the results of studies of pain perception processes using EEG recordings with the model of the ascending pain system(three-route model of pain perception).

This article is organized in three parts. In the first part (sections II-III), we describe the problems of using EEG as a research tool for pain brain activity and propose a hypothetical model of the relationship between brain regions during pain perception. In the second part (sections IV-V), we list the studies used to conduct the construction of the hypothesized model and describe the method of its construction. In the third and final part of the article (section VI) we describe the obtained model of pain perception and compare the results obtained on the basis of literature data.

## 2 Studies of EEG recordings

Studies of pain perception using EEG have been described in detail in systematic reviews[13, 14]. The authors of these reviews come to similar conclusions about the impossibility at the moment to identify brain regions responsible for pain activity using EEG, and as a consequence, the impossibility to use EEG recordings as biomarkers of pain activity.

The authors emphasize the inconsistencies of the results presented in the articles and provide recommendations for future research on this topic. They suggest developing detailed research protocols that would take into account various factors affecting EEG recordings (open/closed eyes, attention disorders during the experiment, etc.); applying machine learning algorithms for data processing; taking into account quantitative characteristics of pain stimulus sources; developing more universal methods of subjective assessment of perceived pain.

The authors of the reviews also propose to further consider the process of pain sensation not as a result of brain activity of one specific brain zone, but as a result of the work of the system of several brain zones.

## 3 Three-pathway model of pain perception

Pain is defined as an unpleasant sensory and emotional experience resulting from actual or potential tissue damage. Central pain processing is known to be carried out by a neural network involving several cortical and thalamic regions of the brain[15]. In healthy control subjects, studies have shown that using brain imaging techniques such as functional magnetic resonance imaging (fMRI), neurons correlate with the applied exogenous nociceptive stimulus[16, 17, 18, 19, 20]. This so-called “pain matrix” includes the thalamus, anterior and posterior insular cortex (aIC and pIC), lateral and medial prefrontal cortex (lPFC and mPFC), anterior, middle and posterior cingulate cortex (ACC, MCC, PCC), primary and secondary somatosensory cortex (S1 and S2), orbitofrontal cortex (OFC), basal ganglia, premotor cortex, midbrain, cerebellum, and posterior parietal cortex (PPC).

A similar scheme of the pain perception process was proposed in the articles[21, 22] based on electrophysiological, anatomical, and psychological studies[16, 23, 24, 25,26, 27, 28, 29]. In this study, the authors examined in detail the influence of emotional and thought processes and attention processes on pain perception. Based on their research, they proposed possible routes by which these processes occur in the brain at the time of pain perception. In this model, the system of pain perception, in which processing is carried out along three different routes, includes the following brain regions: thalamus, insula, cerebellum, anterior cingulate cortex, primary somatosensory cortex, secondary somatosensory cortex, amygdala, basal ganglia, periaqueductal gray, parabrachial nucleus, prefrontal cortex.

According to this scheme, the zones of the anterior cingulate cortex and amygdala at the moment of pain sensation are responsible for processes related to motivation and emotions; S1 and S2 are responsible for decoding information about the place of pain sensation and its duration. Based on researches[30, 31, 32], the area of the prefrontal cortex is responsible for cognitive processes (attention, memory, decision making).

In a study of the insula, S1 and thalamus were activated when pain sensation was presented during imagined movement of the amputated arm.

## 4 Articles for analysis

### 4.1 Criteria for selection of articles

For this analysis, articles were searched using keywords and combinations of “neuropathic pain, nociceptive pain, EEG, thalamus, thalamic activity, amygdala, somatosensory cortex” in PubMed, Scopus, and Google Scholar databases and article libraries. The articles then specified the following criteria for analysis:

1. Articles that investigated nociceptive stimulation of healthy patients should have described in detail the protocols for conducting experiments and algorithms for processing the results. This makes it possible to ensure the reliability and accuracy of the obtained data;
2. The articles should have explicitly stated changes in electroencephalogram (EEG) rhythms. The EEG is a useful tool for studying the electrical activity of the brain and can be used to assess changes associated with nociceptive or neuropathic pain;
3. The articles should also have specified the topological distributions of EEG activity. Topographic analysis allows us to determine which areas of the brain are activated or deactivated in association with nociceptive or neuropathic pain. This helps to better understand which brain regions are associated with pain perception.

Using these criteria, the most relevant research literature was selected for this analysis. This provided a robust and informative set of articles for the study of nociseptive pain and its relationship to changes in EEG rhythms and topological distribution of brain activity.

### 4.2 Articles investigating EEG recordings with nociceptive stimulation

This group included articles that investigated brain activity induced by stimulated nociseptive pain in healthy subjects. Stimulations were carried out by different methods, which included the following methods:

1. Stimulation of thermal pain induced by contact-thermal thermodrome[50, 54, 39, 41];
2. Stimulation of hypothermia of hands in a bucket of ice water[47, 51, 38];
3. Electrical stimulation[40, 49];
4. Intramuscular injection of hypertonic saline solution[42, 55, 56, 52];
5. Topical application of capsaicin cream[48];
6. Stimulation of pain from pressure tourniquet cuffs[57];

In these studies, pain was stimulated in the hands. The results of these studies include descriptions of the results of EEG recordings (localization of rhythm changes, nature of rhythm changes) during these stimulations, images of topographic distributions of EEG rhythm power changes.

### 4.3 Articles on the study of EEG recordings with neuropathic pain

This group included articles investigating neuropathic pain in patients with various neurologic diseases (multiple sclerosis, aging-induced thalamic disorders) using EEG. Studies have observed the following types of neuropathic pain:

1. Central neuropathic pain[53, 43] - these articles dealt with neuropathic pain caused by central nervous system disorders (multiple sclerosis) and spinal cord injuries;
2. Peripheral pain[44, 46] - these articles considered patients with chronic ortofacial neuropathic pain and postherpetic neuralgia;
3. Mixed neuropathic pain [58, 45].

## 5 Method of analysis

For analysis the brain activity recorded by the EEG, there were selected zones of scalp surface, that corresponded to the brain zones proposed in the theory of ascending pain.

### Primary somatosensory cortex

The primary somatosensory cortex (S1) is a well-defined area of the brain that is involved in processing somatosensory information. Electrical activity generated in S1 can be detected on EEG recordings in the central scalp areas (C3 and C4).

### Prefrontal cortex

The prefrontal cortex is an area of the brain that plays a crucial role in cognitive decision-making and working memory. Electrodes associated with the prefrontal cortex on EEG recordings include the frontal area electrodes (Fp1, Fp2, F3, F4, F7, F8, and AFz).

#### The internal structures of the brain

Since a number of the brain regions considered in this model paper are located deep within the brain and cannot be explicitly recorded by EEG, brain surface areas have been identified by which the activity of deep regions can be indirectly ascertained.

### Thalamus

Activity caused by thalamocortical interactions at the brain surface can be captured by changes in alpha and delta rhythm across the brain surface. Anterior electrodes Fp1 and Fp2 showing delta activity and occipital electrodes O1 and O2 showing alpha activity indirectly indicate thalamic activity.

### Insular gyrus

Due to the special position of the islet gyrus relative to the brain surface, it is difficult to record activity in this area using EEG electrodes. Indirectly, the activity of this area can be identified by changes in electrical activity at the frontal (Fz, FCz), central (Cz) and temporal electrodes (T3, T4).

### Amygdala

In studies of processes associated with the amygdala using EEG, changes have been observed at frontal(Fp1, Fp2, F3, F4, F7 and F8) and temporal electrode sites(T7 and T8), making these areas of the cerebral surface possible indicators of amygdala activity[35, 36].

### Secondary somatosensory cortex

Due to the location of S2 and its relationship with S1 on EEG recordings, S2 activity is usually observed in the parietal regions of the skull. The electrodes reflecting these activities are electrodes Pz and CPz[34]. Also, activity in the areas of electrodes Cp3 and Cp4 is also associated with activity of the secondary sensorimotor cortex.

### Basal ganglia

Brain surface activity evoked by the basal ganglia is expressed in the activity of fronto-central (Fz, FCz) and fronto-right (F4) electrodes. These cortical areas are functionally connected to the basal ganglia and may show altered activity in certain motor disorders or conditions affecting the basal ganglia.

### Periaqueductal gray, Parabrachial nucleus, Anterior cingulate cortex

In activity studies, the Periaqueductal gray, Parabrachial nucleus and Anterior cingulate cortex are often examined using the medial frontal(Fz) and central areas(Cz, Fcz).

### Cerebellum

This section of the brain is located deep in the back of the brain, making detection of its activity using electrodes on the surface of the head quite problematic due to the limited spatial resolution of the EEG.

The general scheme of the zones reflecting the activity of the brain areas considered in the article are shown in Fig. 2.

**Figure 1:**
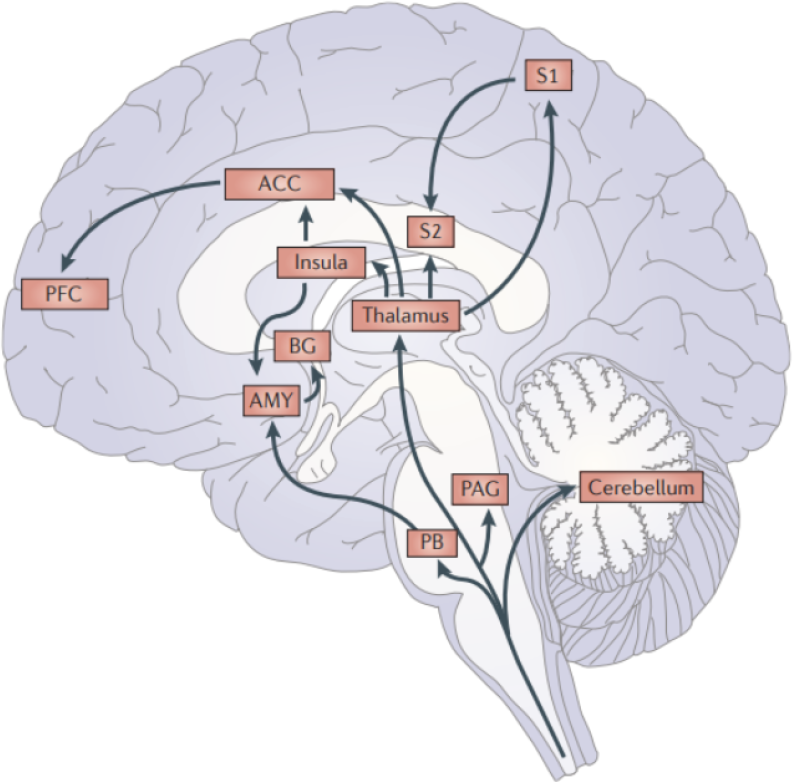
Afferent pain pathways include multiple brain regions: thalamus, insula, cerebellum, anterior cingulate cortex (ACC), primary somatosensory cortex (S1), secondary somatosensory cortex (S2), amygdala (AMY), basal ganglia (BG), periaqueductal gray (PAG), parabrachial nucleus(PB), pre-frontal cortex(PFC)[21].

**Figure 2:**
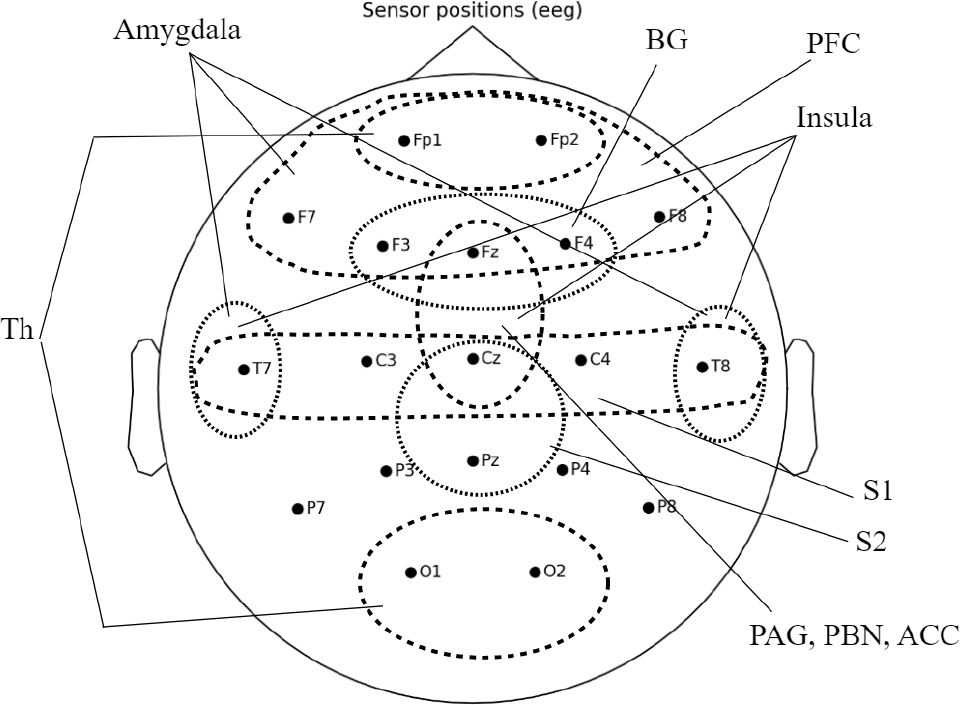
General scheme of the zones reflecting the activity of the brain areas: thalamus(Th), insula, anterior cingulate cortex (ACC), primary so-matosensory cortex (S1), secondary somatosensory cortex (S2), amygdala, basal ganglia (BG), periaqueductal gray (PAG), parabrachial nucleus(PB), prefrontal cortex(PFC)

## 6 Results

The EEG rhythm power changes described in the articles were sorted out by brain areas where these changes were observed. The general results of analysis are shown in Table 1. The averaged across articles spatial distributions are schematized in Figures 3 - 7.

**Table 1:** General results of article analysis:P.F.C-Prefrontal cortex, F.C.-Frontal cortex, P.C.-Parietal cortex, T.C. - Temporal cortex, O.C.-occipital cortex; inc. - increased, dec. - decreased.

| Lobe | Delta | Theta | Alpha | Beta | Gamma |
| --- | --- | --- | --- | --- | --- |
| P.F.C | N.A. | inc. | inc. | inc. | inc. |
| F.C. | inc. | inc./dec. | inc./dec. | inc. | inc. |
| P.C. | inc. | N.A. | inc./dec. | inc. | dec. |
| T.C. | inc. | inc./dec. | dec. | inc. | inc. |
| O.C. | inc. | N.A. | inc./dec. | N.A. | N.A. |

**Figure 3:**
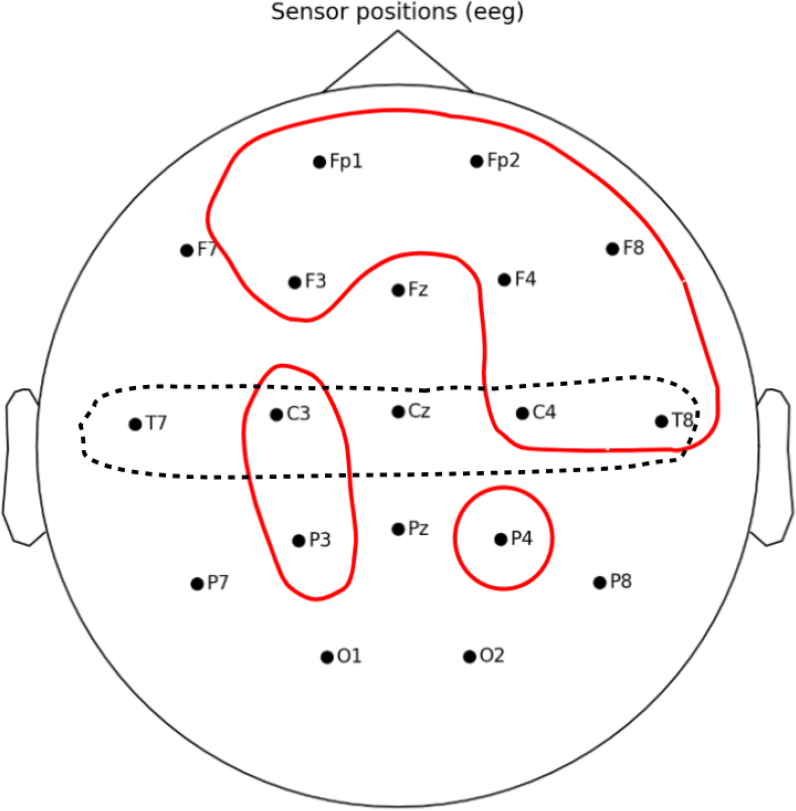
The averaged across articles spatial distribution of power changes of the delta EEG rhythm, red contours are regions, where power of rhythm has increased.The dotted line indicates the area of somatosensory cortex.

**Figure 4:**
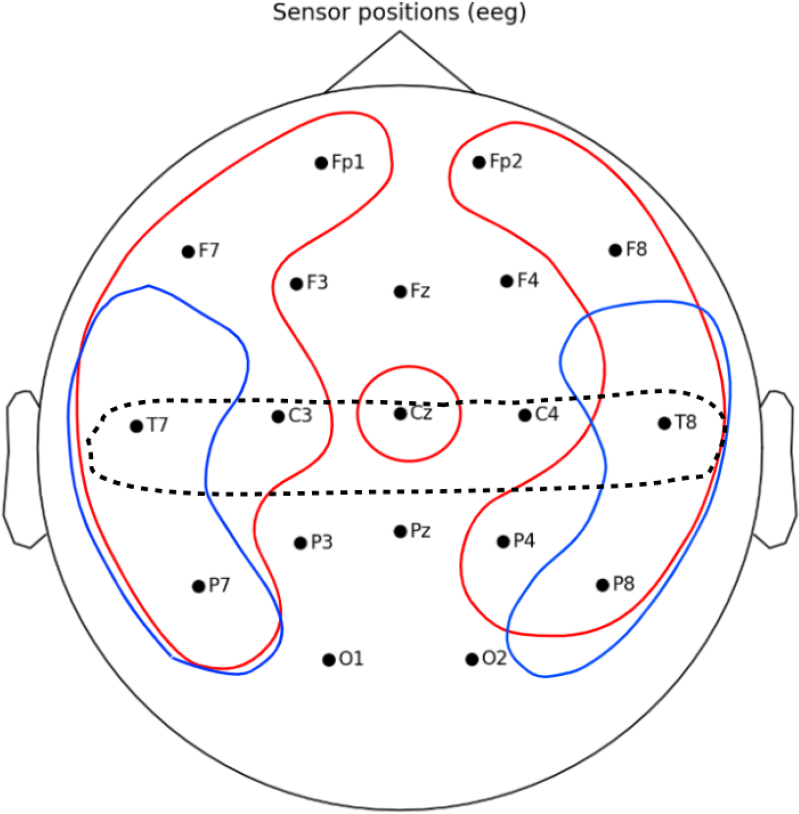
The averaged across articles spatial distribution of power changes of the theta EEG rhythm; red contours are regions, where power of rhythm has increased; blue contours are regions, where power of rhythm has decreased. The dotted line indicates the area of somatosensory cortex.

**Figure 5:**
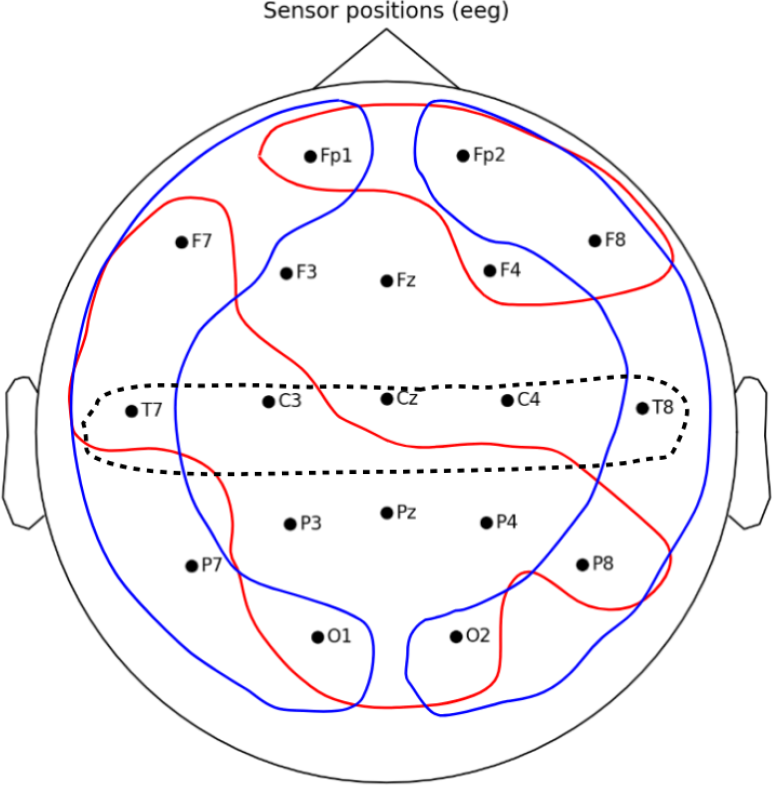
The averaged across articles spatial distribution of power changes of the alpha EEG rhythm; red contours are regions, where power of rhythm has increased; blue contours are regions, where power of rhythm has decreased. The dotted line indicates the area of somatosensory cortex.

**Figure 6:**
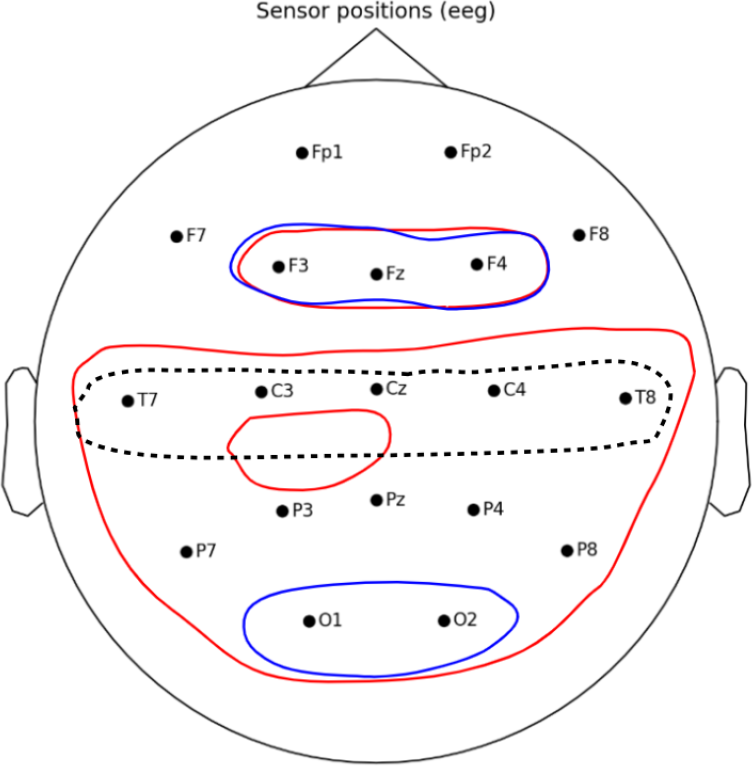
The averaged across articles spatial distribution of power changes of the beta EEG rhythm; red contours are regions, where power of rhythm has increased; blue contours are regions, where power of rhythm has decreased. The dotted line indicates the area of somatosensory cortex.

**Figure 7:**
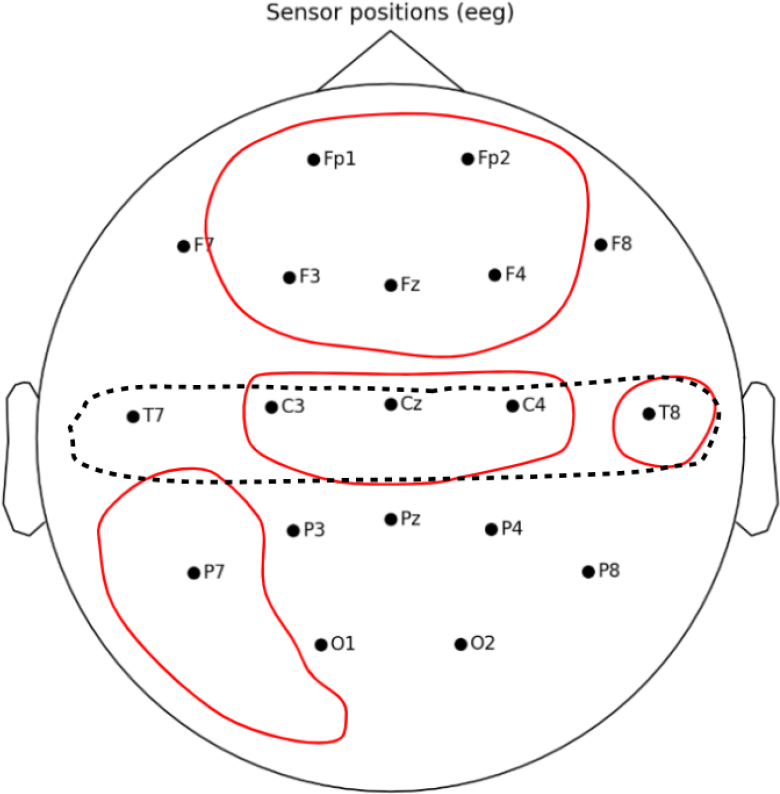
The averaged across articles spatial distribution of power changes of the gamma EEG rhythm; red contours are regions, where power of rhythm has increased; blue contours are regions, where power of rhythm has decreased. The dotted line indicates the area of somatosensory cortex.

### Delta rhythm

In articles during painful stimulations, an increase in the power of delta rhythm was recorded. The distribution of delta rhythm changes is presented in Figure 3.

In the prefrontal lobe of the brain, an increase in delta rhythm was observed in the articles[38, 50]. Increases in delta rhythms in this area were also recorded in the article[40].

Increases in delta rhythm power in the frontal lobe were observed during the experiments in the articles[47, 48]. Increases in delta rhythm power were also observed in frontal lobe areas contralateral to the stimulated hemisphere in the article [42].

The temporal lobes showed an increase in delta rhythm power in the right hemisphere[51].

No changes in delta rhythm were observed in the parietal and occipital lobes.

### Theta rhythm

Changes in the theta rhythm are of mixed character. In frontal, central, temporal and occipital regions, an increase in theta-rhythm power was recorded. At the same time, mixed results were observed in the same temporoparietal regions. The spatial distribution of theta-rhythm changes is presented in Figure 4.

In the prefrontal lobe, an increase in the theta rhythm was observed as described in [40]. A simultaneous increase in theta rhythm power was observed in the medial prefrontal cortex and dorsolateral prefrontal cortex regions in the article[44].

In the frontal cortex, increases in theta power were observed in the bilateral insular cortex and in the contralateral precuneus and posterior cingulate cortex[49]. Increases in theta power were also observed in patients with thalamic nuclei dysfunction in the article[58], and this article also described gradual decreases in theta power after nucleus surgery.

In the temporal lobes, there were significant decreases in the power of the theta rhythm in the left temporal region at the T7 electrode, with a simultaneous increase in the power in the right temporal region at the T8 electrode [49, 50]. In patients with multiple sclerosis, there was an increase in theta rhythm power in the temporal lobes of the right hemisphere[53]. Also, an increase in theta rhythm power during pain stimulation was observed at temporal electrodes in the right hemisphere[51]. Decreases in the theta rhythm were observed in the article[38]. Decreases in theta rhythm in the parietal lobe were described in the article[38].

Patients before surgery on the thalamic nuclei showed increased power of the theta rhythm in the left hemisphere, and as a consequence, a decrease was observed in the article[45] after surgery.

No changes in the theta rhythm were observed in the occipital lobe.

### Alpha rhythm

The spatial distribution of alpha-rhythm changes is presented in Figure 5.

In article[44], an increase in alpha rhythm power was observed in regions of the medial prefrontal cortex and dorsolateral prefrontal cortex.

Increases in alpha rhythm power were observed in the frontal region and correlated with changes in pain stimulus intensity and pain severity[43]. At the same time, decreases in alpha-1 rhythm power in the frontal brain region were observed in the frontal brain region in the articles[50, 38].

Decreases in alpha rhythm power were observed at temporal electrodes of both hemispheres[54]. Decreased alpha-1 activity in temporal brain regions[50, 38].

In the parietal lobe, a decrease in alpha power was observed in most articles[42, 38, 41]. Also, a decrease in power was observed in areas contralateral (P3-P4) to the stimulated arm[47, 48, 56, 57]. Despite this, an increase in alpha rhythm in both parietal regions was described in the paper[55]. Increased alpha rhythm power in the occipital and parietal regions was recorded in the [45].

In the occipital lobe area, both decreases in alpha rhythm in the articles[42, 56, 57] and increases in alpha-1 power in the occipital area were observed in the article[57].

### Beta rhythm

In most of the article, an increase in beta rhythm power was recorded during the sensation of pain. The spatial distribution of theta-rhythm changes is presented in Figure 6.

In article[44], simultaneous increases in beta rhythm power were observed in regions of medial prefrontal cortex and dorsolateral prefrontal cortex.

In the frontal lobe, an increase in beta rhythm power was observed in the frontal lobe during the experiments in the articles[47, 55] and were asymmetric in nature.

An increase in beta rhythm power was recorded in the temporal lobes during the experiments in[42, 44, 47] articles[42, 44, 47]. Increases in beta rhythm power during nociceptive pain studies were observed in contralateral brain regions relative to the stimulated arm(T5,T9)[50, 51]. Increased power of beta-1 and beta-2 rhythms was observed in patients with multiple sclerosis and with neuropathic pain in the temporal lobes of both hemispheres[53].

In the articles[51, 39] in bilateral posterior parietal lobe regions, an increase in beta-1 rhythm power was described. An increase in beta-2 rhythm in the right posterior parietal region was described in the article[55, 38]. A decrease in beta rhythm was observed in the article[41].

In the occipital regions of both hemispheres, an increase in beta rhythm power was recorded in[48, 53, 45].

### Gamma rhythm

The articles described the increase in gamma rhythm power. The distributions of gamma rhythm changes are presented in Figure 7.

Increases in gamma rhythm power were recorded in the prefrontal area during pain[42, 39, 41, 46].

An increase in gamma rhythm power in frontal central and central frontal lobe regions during painful stimuli was observed in the articles[42, 52]. Also during painful stimuli, a significant increase in gamma rhythm power was observed in the contralateral to handedness brain region in frontal brain areas(FT7), while a decrease in gamma rhythm power was observed in the ipsilateral region(FT8)[49]. Increases in gamma rhythm power in temporal lobes were observed in[46].

In the parietal zones, an increase in the magnitude of the gamma rhythm was observed in the article[38].

No changes in gamma rhythm were observed in the occipital and temporal lobes.

## 7 Discussion

The studies presented in this analysis demonstrate that during the process of pain perception, activity on the surface of the brain changes throughout the brain, supporting the hypothesis that several connected sections of the brain, rather than a specific brain region, are responsible for pain perception in the brain.

Characteristic changes in EEG rhythms can be distinguished for each zone:

- In **the prefrontal cortex**, increases in the power of delta, theta, and gamma rhythms were observed. The alpha rhythm in this area showed mixed power changes. Beta rhythm almost did not change;
- **The primary somatosensory cortex** showed an increase in beta and gamma rhythm power. Theta and alpha rhythms showed mixed results. The delta rhythm was virtually unchanged in this area. Also in this area the rhythms changed contralaterally to the stimulated body parts;
- Changes in delta rhythm in areas Fp1 and Fp2 and changes in alpha rhythm in occipital cortex indicate activity of **the thalamus** in pain perception processes;
- The following changes were observed in the areas of frontal and temporal electrodes indirectly indicating the activity of **the insular gyrus**. In the temporal lobes mixed changes of theta and alpha rhythm power were observed. Only in the right temporal lobe there were increases in delta and gamma rhythms. Beta rhythm increased in all areas responsible for the insular gyrus;
- In frontal and temporal regions responsible for the activity of **the amygdala**, there was an increase in the power of delta, theta, and gamma rhythms. The alpha rhythm both increased and decreased in these areas. Beta rhythm showed mixed changes only in frontal electrodes(F3, Fz, F4);
- In the area reflecting the activity of **periaqueductal gray, parabrachial nucleus** and **anterior cingulate cortex**, delta and alpha rhythms did not change, theta rhythm increased at the central electrode(Cz), beta and gamma rhythms increased in the whole area;
- In the **secondary somatosensory cortex** region, delta and gamma rhythms did not change, theta rhythm increased only in the right region, which could be due to a contralateral sensory response, and alpha and beta rhythms increased;
- In the area reflecting **basal ganglia** activity, there was an increase in delta rhythm only around electrode F4, theta and alpha rhythms did not change, beta rhythm showed mixed results, gamma rhythm increased over the whole area.

Increased activity in the prefrontal cortex correlated with the processes of higher nervous activity during the experiments - attention, learning, planning of executive functions, which determines the role of this section of the brain in the processing of pain processes. In the parietal lobe of the brain in the area of somatosensory cortex there were clearly recorded synchronous with changes in pain stimuli decreases in alpha rhythm power, which indicates the role of this section of the brain in pain perception - spatial perception of pain in the body.

Despite research, the use of EEG to record brain activity associated with pain is complicated by spatial resolution and the complex structure of the brain - many areas responsible for pain perception are located under the outer cortex, which allows only indirectly recording their activity. As a result, in order to obtain more detailed information for studying this activity, it is necessary to use additional non-invasive methods of brain research together with EEG: fMRI, MEG.

One possible solution to increase the accuracy of pain activity studies may be the application of machine learning algorithms to the study of EEG recordings.

An important aspect in the issue of research on brain activity of pain sensations is the aspect of interconnection of different brain regions and the influence of their interconnections on the observed activity - coherence of brain sections. Such a study was cited in the article[51], which analyzed the coherence of changes in the rhythms of different parts of the brain and their influence on neighboring areas. The use of machine learning algorithms to investigate this aspect of the pain perception process may increase the accuracy of the research.

## References

[1] MSD Manual Professional Edition. (Mar 2022). Overview of Pain[Online]. Available: https://www.msdmanuals.com/professional/neurologic-disorders/pain/overview-of-pain

[2] K. H. Brodersen, K. Wiech, E. I. Lomakina, C. S. Lin, J. M. Buhmann, U. Bingel, M. Ploner, K. E. Stephan, I. Tracey, et al., “Decoding the perception of pain from fMRI using multivariate pattern analysis,” NeuroImage, vol. 63, pp. 1162–1170, Nov 2012.

[3] S. E. Longe, R. Wise, S. Bantick, D. Lloyd, H. Johansen-Berg, F. McGlone, I Tracey, et al., “Counter-stimulatory effects on pain perception and processing are significantly altered by attention: an fMRI study,” Neuroreport, vol. 12, pp. 2021–2025, Jul 2001.

[4] M. Damascelli. T. S. Woodward, N. Sanford, H. B. Zahid, R. Lim, A. Scott, J. K. Kramer, et al., “Multiple Functional Brain Networks Related to Pain Perception Revealed by fMRI,” Neuroinformatics, vol. 20, pp. 155–172. Jan 2022.

[5] K. Hugdahl, G. Rosén, L. Ersland, A. Lundervold, A. I. Smievoll, R. Barndon, T. Thomsen, et al., “Common pathways in mental imagery and pain perception: An fMRI study of a subject with an amputated arm,” Scandinavian Journal of Psychology, vol. 42, pp. 269–275, Jul 2001.

[6] G. L. Raff, K. M. Chinnaiyan, R. C. Cury, M. T. Garcia, H. S. Hecht, J. E. Hollander, B. O’Neil, A. J. Taylor, U. Hoffmann, et al., “SCCT guidelines on the use of coronary computed tomographic angiography for patients presenting with acute chest pain to the emergency department: A Report of the Society of Cardiovascular Computed Tomography Guidelines Committee,” Journal of Cardiovascular Computed Tomography, vol. 8, pp. 254–271, Jun 2014.

[7] A. B. Newberg, P. J. LaRiccia, B. Y. Lee, J. T. Farrar, L. Lee, A. Alavi, et al., “Cerebral blood flow effects of pain and acupuncture: a preliminary single-photon emission computed tomography imaging study” Journal Neroimaging, vol. 15, pp. 43–49, Jan 2005.

[8] Y. Kanpolat, H. C. Ugur, M. Ayten, A. H. Elhan, et al., “Computed tomography-guided percutaneous cordotomy for intractable pain in malignancy,” Operative Neurosurgery, vol. 64, pp. 187–93, Mar 2009.

[9] I.-W. Su, F.-W. Wu, K.-C. Liang, K.-Y. Cheng, S.-T. Hsieh, W.-Z. Sun, T.-L. Chou, et al., “Pain Perception Can Be Modulated by Mindfulness Training: A Resting-State fMRI Study,” Frontiers in Human Neuroscience, vol. 10, Nov 2016.

[10] X. Xu, H. Fukuyama, S. Yazawa, T. Mima, T. Hanakawa, Y. Magata, M. Kanda, N. Fujiwara, K. Shindo, T. Nagamine, H. Shibasaki, et al., “Functional localization of pain perception in the human brain studied by PET,” NeuroReport, vol. 8, pp. 555–559, Jan 1997.

[11] M. Boly, M.-E. Faymonville, C. Schnakers, P. Peigneux, B. Lambermont, C. Phillips, P. Lancellotti, A. Luxen, M. Lamy, G. Moonen, P. Maquet, S. Laureys, et al., “Perception of pain in the minimally conscious state with PET activation: an observational study,” Lancet Neurology, vol. 7, pp. 1013–1020, Nov 2008.

[12] C. Muller, A. Klega, H.-G. Buchholz, R. Rolke, W. Magerl, R. Schirrmacher, E. Schirrmacher, F. Birklein, R.-D. Treede, M. Schreckenberger, et al., “Basal opioid receptor binding is associated with differences in sensory perception in healthy human subjects: A [18F]diprenorphine PET study,” NeuroImage, vol. 49, pp. 731–737, Jan 2010

[13] P. Zis, A. Liampas, A. Artemiadis, et al., “EEG recordings as biomarkers of pain perception: where do we stand and where to go?” Pain Therapy, vol. 11, pp. 369–380, Jun 2022.

[14] T. Mussigmann, B. Bardel, J.P. Lefaucheur, et al., “Resting-state electroencephalography (EEG) biomarkers of chronic neuropathic pain. A systematic review,” Neuroimage, vol. 1, pp. 369–380, Jun 2022.

[15] R. Melzack and K. L. Casey, et al., “Sensory, motivational, and central control determinants of pain: a new conceptual model,” In D. Kenshalo (Ed.), The skin senses, pp. 423–443, Sprinfield, IL: Thomas.

[16] A. V. Apkarian, M. C. Bushnell, R.-D. Treede, J.-K. Zubieta, et al., “Human brain mechanisms of pain perception and regulation in health and disease,” European Journal of Pain, vol. 9, pp. 463–463, Aug 2005.

[17] A. C. N. Chen, W. Feng, H. Zhao, Y. Yin, P. Wang, et al., “EEG default mode network in the human brain: Spectral regional field powers,” NeoroImage, vol. 41, pp. 561–574, Jun 2008.

[18] A. A. Dubé, M. Duquette, M. Roy, F. Lepore, G. Duncan, P. Rainville, et al., “Brain activity associated with the electrodermal reactivity to acute heat pain,” NeuroImage, vol. 45, pp. 169–180, Dec 2008.

[19] X. Moisset, D. Bouhassira “Brain imaging of neuropathic pain,” NeuroImage, vol. 37, Feb 2007.

[20] R. Peyron, B. Laurent, L. García-Larrea, et al., “Functional imaging of brain responses to pain. A review and meta-analysis”, Journal of Clinical Neurophysiology, vol. 30, pp. 263–288, Oct 2000.

[21] M. C. Bushnell, M. Ceko, L. A. Low, et al., “Cognitive and emotional control of pain and its disruption in chronic pain,” Nature Reviews Neuroscience, vol. 14, pp. 502–511, Jul 2013.

[22] L. Duque, G. Fricchione, et al., “Fibromyalgia and its New Lessons for Neuropsychiatry,” Medical Science Monitor Basic Research, vol. 25, pp. 169–178, Jul 2019.

[23] A. W. Roe, L. M. Chen, R. Friedman, “Intrinsic Signal Imaging of Somatosensory Function in Nonhuman Primates,” in “The Senses: A Comprehensive Reference,”, 2020, pp. 288–301.

[24] E. Rausell, E. G. Jones, et al. “Histochemical and immunocytochemical compartments of the thalamic VPM nucleus in monkeys and their relationship to the representational map,” Journal of Neuroscience, vol. 11, pp. 210–225, Jan 1991.

[25] A. V. Apkarian, T. Shi, et al. “Viscerosomatic interactions in the thalamic ventral posterolateral nucleus (VPL) of the squirrel monkey,” Brain Research, vol.787, pp. 269–276, Mar 1998.

[26] A. D. Craig, J. O. Dostrovsky, et al., “Thermoreceptive lamina I trigeminothalamic neurons project to the nucleus submedius in the cat,” Experimental Brain Research, vol. 85, pp. 470–474, Jun 1991.

[27] R. P. Dum, D. J. Levinthal, P. L. Strick, et al., “The spinothalamic system targets motor and sensory areas in the cerebral cortex of monkeys,” Journal of Neuroscience, vol. 29, pp. 14223–14235, Nov 2009.

[28] C. Y. Saab, W. D. Willis, et al., “The cerebellum: organization, functions and its role in nociception,” Brain Research Reviews, vol. 42, pp. 85–95, Apr 2003.

[29] L. Monconduit, L. Villanueva,et al., “The lateral ventromedial thalamic nucleus spreads nociceptive signals from the whole body surface to layer I of the frontal cortex,” European Journal of Neuroscience, vol. 21, pp. 3395–3402, Jul 2005.

[30] N. P. Friedman, T. W. Robbins, et al., “The role of prefrontal cortex in cognitive control and executive function,” Neuropsychopharmacology, vol. 47, pp. 72–89, Jan 2022.

[31] S. V. Siddiqui, U. Chatterjee, D. Kumar, A. Siddiqui, N. Goyal, et al., “Neuropsychology of prefrontal cortex,” Indian Journal of Psychiatry, vol. 50, pp. 202–208, Jul 2008.

[32] C. Frith, R. Dolan, et al., “The role of the prefrontal cortex in higher cognitive functions,” Brain research. Cognitive brain research, vol. 5, pp. 175–181, Dec 1996.

[33] G. Rosen, F. Willoch, P. Bartenstein, N. Berner, S. Rosjo, et al., “Neurophysiological processes underlying the phantom limb pain experience and the use of hypnosis in its clinical management: an intensive examination of two patients,” International Journal of Clinical and Experimental Hypnosis, vol. 49, pp. 38–55, Feb 2001.

[34] A. Melnik, W. D. Hairston, D. P. Ferris, P. König, et al., “EEG correlates of sensorimotor processing: independent components involved in sensory and motor processing,” Scientific Reports, vol. 7,Jun 2017.

[35] S. Sonkusare, D. Qiong, Y. Zhao, W. Liu, R. Yang, A. Mandali, L. Manssuer, C. Zhang, C. Cao, B. Sun, S. Zhan, V. Voon, et al., “Frequency dependent emotion differentiation and directional coupling in amygdala, orbitofrontal and medial prefrontal cortex network with intracranial recordings,” Molecular Psychiatry, vol. 28, pp.1636–1646, Dec 2022.

[36] V. Zotev, H. Yuan, M. Misaki, R. Phillips, K. D. Young, M. T. Feldner, J. Bodurka, et al., “Correlation between amygdala BOLD activity and frontal EEG asymmetry during real-time fMRI neurofeedback training in patients with depression,” NeuroImage: Clinical, vol. 11, pp. 224–238, Feb 2016.

[37] P. Li, W. Peng, H. Li, C. B. Holroyd, et al., “Electrophysiological measures reveal the role of anterior cingulate cortex in learning from unreliable feedback,” Cognitive, Affective, and Behavioral Neuroscience, vol.18, pp. 949–963, Oct 2018.

[38] M. Gram, C. Graversen, S. S. Olesen, A. M. Drewes, et al., “Dynamic spectral indices of the electroencephalogram provide new insights into tonic pain,” Clinical Neurophysiology, vol. 126, pp. 763–771, Apr 2015.

[39] E. Schulz, E. S. May, M. Postorino, L. Tiemann, M. M. Nickel, V. Witkovsky, P. Schmidt, J. Gross, M. Ploner, et al., “Prefrontal Gamma Oscillations Encode Tonic Pain in Humans,” Cerebral Cortex, vol. 25, pp. 4407–4414, Nov 2015.

[40] R. D. Rouleau, L. Lagrandeur, K. Daigle, D. Lorrain, G. Léonard, K. Whittingstall, P. Goffaux, et al., “Significance of Non-phase Locked Oscillatory Brain Activity in Response to Noxious Stimuli,” Canadian Journal of Neurological Sciences, vol. 42, pp. 436–443, Nov 2015.

[41] S. F. Bunk, S. Lautenbacher, J. Rüsseler, K. Müller, J. Schultz, M. Kunz, et al., “Does EEG activity during painful stimulation mirror more closely the noxious stimulus intensity or the subjective pain sensation?” Somatosensory and Motor Research, vol. 35, pp.192–198, Dec 2018.

[42] P. Veerasarn, C. S. Stohler, et al., “The effect of experimental muscle pain on the background electrical brain activity,” PAIN, vol. 49, pp. 349–360, Jun 1992.

[43] M. P. Jensen, L. H. Sherlin, K. J. Gertz, A. L. Braden, A. E. Kupper, A. Gianas, J. D. Howe, S. Hakimian, et al., “Brain EEG activity correlates of chronic pain in persons with spinal cord injury: clinical implications,” Spinal Cord, vol. 51, pp. 55–58, Jan 2013.

[44] F. Di Pietro, P. M. Macey, C. D. Rae, Z. Alshelh, V. G. Macefield, E. R. Vickers, L. A. Henderson, et al., “The relationship between thalamic GABA content and resting cortical rhythm in neuropathic pain,” Hum Brain Mapping, vol. 39, pp. 1945–1956, May 2018.

[45] L. Michels, M. Moazami-Goudarzi, D. Jeanmonod, et al.,”Correlations between EEG and clinical outcome in chronic neuropathic pain: surgical effects and treatment resistance,” Brain Imaging and Behavior, vol. 5, pp. 329–348, Sep 2011.

[46] R. Zhou, J. Wang, W. Qi, F.-Y. Liu, M. Yi, H. Guo, Y. Wan, et al.,”Elevated Resting State Gamma Oscillatory Activities in Electroencephalogram of Patients With Post-herpetic Neuralgia,” Frontiers in Neuroscience, vol. 12, p. 750, Oct 2018.

[47] S. Ferracuti, S. Seri, D. Mattia, G. Cruccu, et al., “Quantitative EEG modifications during the Cold Water Pressor Test: Hemispheric and hand differences,” International Journal of Psychophysiology, vol. 17, pp. 261–268, Aug 1994.

[48] P.F. Chang, L. Arendt-Nielsen, T. Graven-Nielsen, P. Svensson, A. C. Chen, et al., “Different EEG topographic effects of painful and non-painful intramuscular stimulation in man,” Experimental Brain Research, vol. 141, pp. 195–203, Nov 2001.

[49] P. Taesler, M. Rose, et al., “Prestimulus theta oscillations and connectivity modulate pain perception,” Journal of Neuroscience, vol. 36, pp. 5026–5033, May 2016.

[50] M. T. Huber, J. Bartling, D. Pachur,S. V. Woikowsky-Biedau, S. Lautenbacher, et al., “EEG responses to tonic heat pain,” Experimental Brain Research, vol. 173, pp. 14–24, Aug 2006.

[51] A.C. Chen, P. Rappelsberger, O. Filz, et al.,”Topology of EEG coherence changes may reflect differential neural network activation in cold and pain perception,” Brain Topography, vol. 11, pp. 125–132, 1998.

[52] L. Li, X. Liu, C. Cai, et al.,”Changes of gamma-band oscillatory activity to tonic muscle pain,” Neuroscience Letters, vol. 627, pp. 126–131, Aug 2016.

[53] N. A. Krupina, M. V. Churyukanov, M. L. Kukushkin, N. N. Yakhno, et al., “Central Neuropathic Pain and Profiles of Quantitative Electroencephalography in Multiple Sclerosis Patients,” Frontiers in Neurology, vol. 10, p. 1380, Jan 2020.

[54] R. R. Nir, A. Sinai, R. Moont, E. Harari, D. Yarnitsky, et al., “Tonic pain and continuous EEG: prediction of subjective pain perception by alpha-1 power during stimulation and at rest,” Clinical Neurophysiology, vol. 123, pp. 605–612, Mar 2012.

[55] D. Le Pera, P. Svensson, M. Valeriani, I. Watanabe, L. Arendt-Nielsen, A. C. Chen, et al., “Long-lasting effect evoked by tonic muscle pain on parietal EEG activity in humans,” Clinical Neurophysiology, vol. 111, pp. 2130–2137, Dec 2000.

[56] P. F. Chang, L. Arendt-Nielsen, T. Graven-Nielsen, A. C. Chen, et al., “Psychophysical and EEG responses to repeated experimental muscle pain in humans: pain intensity encodes EEG activity,” Brain Research Bulletin, vol. 59, pp. 533–543, Feb 2003.

[57] L. L. Egsgaard, L. Wang, L. Arendt-Nielsen, et al., “Volunteers with high versus low alpha EEG have different painEEG relationship: a human experimental study,” Experimental Brain Research, vol. 193, pp. 361–369, Mar 2009.

[58] J. Sarnthein, J. Stern, C. Aufenberg, V. Rousson, D. Jeanmonod, et al., “Increased EEG power and slowed dominant frequency in patients with neurogenic pain,” Brain, vol. 129, pp. 55–64, Jan 2006.

